# Effects of different feeding methods on serum biochemical indexes, metabolic indexes, immune indexes, and intestinal microorganisms of Nanjiang yellow goats

**DOI:** 10.1101/2024.01.10.575136

**Authors:** Yangyang Luo, Asma Anwar, Cheng Pan, Hengbo Shi, Shehr Bano Mustafa, Yu Chen, Zhenzhen Zhang, Jingjing Li, Jiangjiang Zhu, Wangsheng Zhao

**Affiliations:** College of Life Sciences and Engineering, Southwest University of Science and Technology, Mianyang, Sichuan 621000, China; Institute of Dairy Science, College of Animal Sciences, Zhejiang University, Hangzhou 310058, China; Sichuan Nanjiang Yellow Goats Original Breeding Farm, Nanjiang, Sichuan 636600, China; Qinghai-Tibetan Plateau Animal Genetic Resource Reservation and Utilization Key Laboratory of Sichuan Province, Chengdu, Sichuan, 610041, China; Key Laboratory of Qinghai-Tibetan Plateau Animal Genetic Resource Reservation and Utilization (Southwest Minzu University), Ministry of Education, Chengdu, Sichuan, 610041, China

**Keywords:** Nanjiang yellow goats, different feeding methods, gut microorganisms, immune indexes, metabolome

## Abstract

The intestinal microbiota significantly influences the intake, storage, and absorption of nutrients in animals, thereby greatly impacting the growth and development of the animals. Factors such as diet, animal breed, growth stage, and feeding methods may affect variations in the composition of the intestinal microbiota. However, research on the variations in the intestinal microbiota and metabolites of Nanjiang Yellow goats under different feeding methods is still unclear. We measured various serum biochemical indicators and immune biochemical indicators and found that the triglyceride (TC) content in the grazing group (the FMMF) was significantly lower than in the barn-feeding group (the SSMF) (*P*<0.05). Serum levels of immunoglobulin A (IgA), immunoglobulin G (IgG), and immunoglobulin G (IgM) were higher in the FMMF group. At the phylum level, the most abundant bacteria were *Firmicutes, Bacteroidota*, and *Verrucomicrobiota*. At the genus level, the most abundant microbial groups were *Christensenellaceae_R-7_group, UCG-005*, and *Rikenellaceae_RC9_gut_group*. Differential metabolite enrichment analysis through KEGG pathways revealed that the most remarkably enriched pathway was “Metabolic pathways,” including Steroid hormone biosynthesis and Arachidonic acid metabolism, among others. Analyzed by multi-omics association, we identified notably different microbial features correlated with immune indicators and metabolites after different feeding methods. We observed a significant negative correlation (*P*<0.05) between the concentrations of serum immune factors interleukin-2 (IL-2), interleukin-4 (IL-4), and *g__probable_genus_10*. The concentration of IgM in serum showed a highly significant positive correlation (*P*<0.01) with the relative abundance of *g__Erysipelatoclostridium* in the intestine. Interestingly, most differential metabolites were significantly associated with intestinal microbiota. This experiment indicates that different feeding methods may influence the diversity and relative abundance of the intestinal microbiota in Nanjiang Yellow goats. The intestinal microbiota is correlated with immune indicators and metabolism, and regulating the diversity and relative abundance of the intestinal microbiota can be a way to adjust metabolism, thereby promoting the healthy growth of the Nanjiang Yellow goats.

## 1. Introduction

The Nanjiang Yellow goats are a breed of goats specializing in Nanjiang County, Sichuan, China, and one of China’s nationally protected animal breeds. It is a meat goat, known for its tasty, tender meat and moderate fat content, and it has outstanding production performance, a fast growth rate, and the ability to reach the standard slaughter weight in a relatively short period[1]. It has been found that different feeding modes can affect the composition of intestinal microorganisms, as well as the physiological functions and metabolism of Nanjiang yellow goats. Nowadays, the implementation of relevant policies has brought pressure on the traditional breeding of grazing livestock, and at the same time, grazing is also causing resource destruction as well as environmental pollution, so the grazing feeding mode is gradually eliminated and replaced by centralized animal feeding operations. To solve these problems, it is necessary to accelerate the transformation of the current traditional feeding management, and accelerate the effective promotion of shelter feeding technology and management mode, to lay a good and solid foundation for the high and stable production of goat’s meat. Understanding the diversity of ruminant gut microbial changes and nutrient metabolism mechanisms is an important basis for improving animal welfare, animal productivity, and animal product quality, as well as reducing greenhouse gas emissions through human intervention[2].

Gut microorganisms play important physiological functions and protective roles in the organism. They participate in food digestion and nutrient absorption, help synthesize vitamins and other beneficial substances, regulate the immune system, maintain the intestinal barrier function, and inhibit the growth of pathogenic microorganisms. Gut microorganisms are closely related to the health of the organism, and are associated with intestinal diseases, metabolic diseases, and immune system diseases. In addition, different feeding modes may have a substantial impact on microbial microbiota composition, which can offer new possibilities for manipulating health through different feeding modes.

Nowadays, it is important to better define the main commensal bacteria, microbiota profiles, and systemic characteristics that produce stable gut microbiota with health benefits. The degree of inter-individual variation in the composition of the microbiota in the animal organism is now also becoming apparent and may influence individual responses to drug administration and dietary manipulation[3]. It has been shown that gut microbial diversity is higher in grazing ruminants than in barn-feeding fee ruminants[4], Zhang et al. found that these results indicate that there are significant differences in the structure and composition of the intestinal microbiome of barn-feeding and grazing Charolais and grazing Charolais have higher coarse fiber digestion ability[5]. Based on previous research, feeding Sunit goats in a confined space improved their weight gain and slaughter performance, but resulted in a reduction in the diversity of microorganisms in their gut, which negatively affected the quality and taste of their meat[6]. Previous research has shown that the way animals are fed can impact their gut microbiota, growth, and immune system. This study focuses on how different feeding methods affect the gut metabolism and function of Nanjiang yellow goats. The aim is to provide a theoretical basis for the optimal feeding culture of this breed and to support efficient breeding practices.

## 2. Materials and methods

### 2.1. Experiments Animal Grouping and Management

The animals used in this experiment were obtained from the Nanjiang Yellow Goats Original Breeding Farm, Dahe Town, Nanjiang County, Bazhong City, Sichuan Province, China, with longitude: 106.74 ° latitude: 31.70 ° elevation at 650-1593.6 m, annual precipitation around 1400-1500 mm, annual average temperature around 12.2 °C (38 °C-8 °C), air relative humidity at 79%, and a frost-free period of roughly 220 days. Twelve Nanjiang yellow goats that had been weaned at the age of two months, with similar body weights and in good health were selected, of which 6 were in the SSFM and 6 were in the FMMF. The SSMF was fed at 7:30 a.m. and 5:30 p.m. every day, while the FMMF was grazed at 7:00 am. every day and backed to the barn at 6:00 pm. The experimental period consisted of a pre-feeding period of 7 days and an orthogonal feeding period of 60 days, totaling 67 days.

### 2.2. Methodology

#### 2.2.1 Sample collection and preservation of Nanjiang yellow goats

The last 5 days of the feeding experiment were used as the blood and feces collection period of the samples (the first 3 days were the physiological adaptation period, and the last 2 days were the fasting sample collection period). The neck blood of each sample goat was collected by a non-enzymatic blood collection tube, and the blood samples collected by each goat were placed in an ice box for temporary preservation. After all the samples were collected, the supernatant was extracted by centrifugation in a 3500 r / min centrifuge at 4 °C for 10 min, which was sub-packed in a 2 mL cryopreserved tube and stored in liquid nitrogen for the determination of serum biochemical indicators. The 12 Nanjiang yellow goats in the SSMF group and the FMMF group were placed in the buttocks with fecal bags and then collected separate fecal samples at 8 : 00,12: 00,16: 00 and 20: 00 every day. The collected feces must be immediately put into carbon dioxide ice for proper preservation. When the sampling period of Nanjiang yellow goats’ feces is over, the collected samples will be mixed evenly. Each feces were collected 15-20 g and above. The mixed fecal samples are sealed and placed in a prepared liquid nitrogen tank to facilitate the immediate return to the laboratory, and placed in a -80 °C ultra-low temperature refrigerator for storage.

#### 2.2.2 Measurement of serum biochemical and immunological indices

The collected serum and rumen fluid were thawed at 4 °C, and the total protein (TP), urea nitrogen (BUN), glucose (GLU), total cholesterol (T-CHO), and TG contents of serum were determined according to the assay method provided by Nanjing Jiancheng reagent kit. The serum levels of IgA, IgG, IgM, IL-2, IL-4, IL-6, and tumor necrosis factor-alpha (TNF-*α*) were determined by Nanjing Jiancheng Elisa kit.

#### 2.2.3 Extraction determination and analysis of non-targeted metabolites

Metabolomics was determined at Hangzhou Lianchuan Biotechnology Co. An aliquot of 200 mg sample was weighed and ground with liquid nitrogen. Lipids were extracted by adding 120 μL of 50% methanol to the sample and mixing well with shaking, and the sample was allowed to stand at room temperature for 10 min. The extract was placed at -20 °C overnight to precipitate the proteins in the sample. The precipitated sample was centrifuged at 4000 g for 20 min, and the supernatant lipid extract was transferred to a 96-well plate. The samples were diluted with diluent (isopropanol: acetonitrile: water = 2:1:1, v/v) to dilute the lipid extract. An aliquot of 10 μL of diluent was removed from each sample and mixed to form a QC sample. All metabolic samples were then stored in the refrigerator at -80 °C before sampling. The raw data of mass spectrometry were converted into readable data mzXML by MSConvert software of Proteowizard. Peak extraction was performed using XCMS software and peak extraction quality control was performed. The extracted substances were subjected to additive ion annotation using CAMERA, and then primary identification was performed using metal software. The first level information of mass spectrometry was used for identification and the second level information of mass spectrometry was used to match with the in-house standard database. Candidate substances were annotated using HMDB, KEGG, and other databases to explain the physicochemical properties and biological functions of the metabolites. The meta software was used for the quantification of differential metabolites and differential metabolite screening.

#### 2.2.4 16S rRNA determination of intestinal microbiota

One copy of the collected and dispensed fecal samples from each goat was sent to Hangzhou Lianchuan Biotechnology Co. Ltd. for sequencing, and the sequencing platform was Illumina Hiseq. The composition and diversity of the intestinal microbiota were determined using 16S rRNA technology. Then the DNA of the gut microbial samples from goats in the SSMF and FMMF groups of Nanjiang yellow goats was extracted using the OMEGA stool DNA kit (Omega Bio-Tek, the U.S). Then the quality of the DNA was assessed using 1.5% agarose gel electrophoresis. The data obtained from the sequencing platform was analyzed for the evaluation of the diversity of the OTUs and the variable statistic isobaric harvesting of the bacterial microbiota structure, in this study, we use the Spearman test method to select the top 30 genera of relative abundance for correlation analysis with metabolic indexes and immune indexes.

### 2.3 Data processing

Using SPSS 26.0 software for statistical analysis and conducting T-tests on the blood biochemical indices of samples, and statistical analysis on the diversity and relative abundance of intestinal microbiota. P < 0.05 was considered as a significant difference and P < 0.01 was considered as a highly significant difference, and the results were expressed as mean and standard error of the mean (SEM). The correlation between gut microbiota and metabolome and the correlation between gut microbiota and immune indicators were also analyzed using Spearman’s test. Finally, GraphPad prism9 software was utilized for plotting.

## 3. Results

### 3.1 Effects of different feeding methods on serum biochemical indexes of Nanjiang Yellow Goats

As shown in **Fig. 1A**, TC content in the FMMF was notably decreased compared to the SSMF (*P*<0.05); serum BUN content in the FMMF was lower than FMMF the SSMF but there was no substantial difference; GLU content in FMMF was lower than the SSMF but there was no significant difference. T-CHO content in the SSMF was higher than that in the SSMF but there was no substantial difference; As shown in **Fig. 1B**, the TP content of the FMMF was higher than the SSMF but without significantly different.

**Fig. 1.**
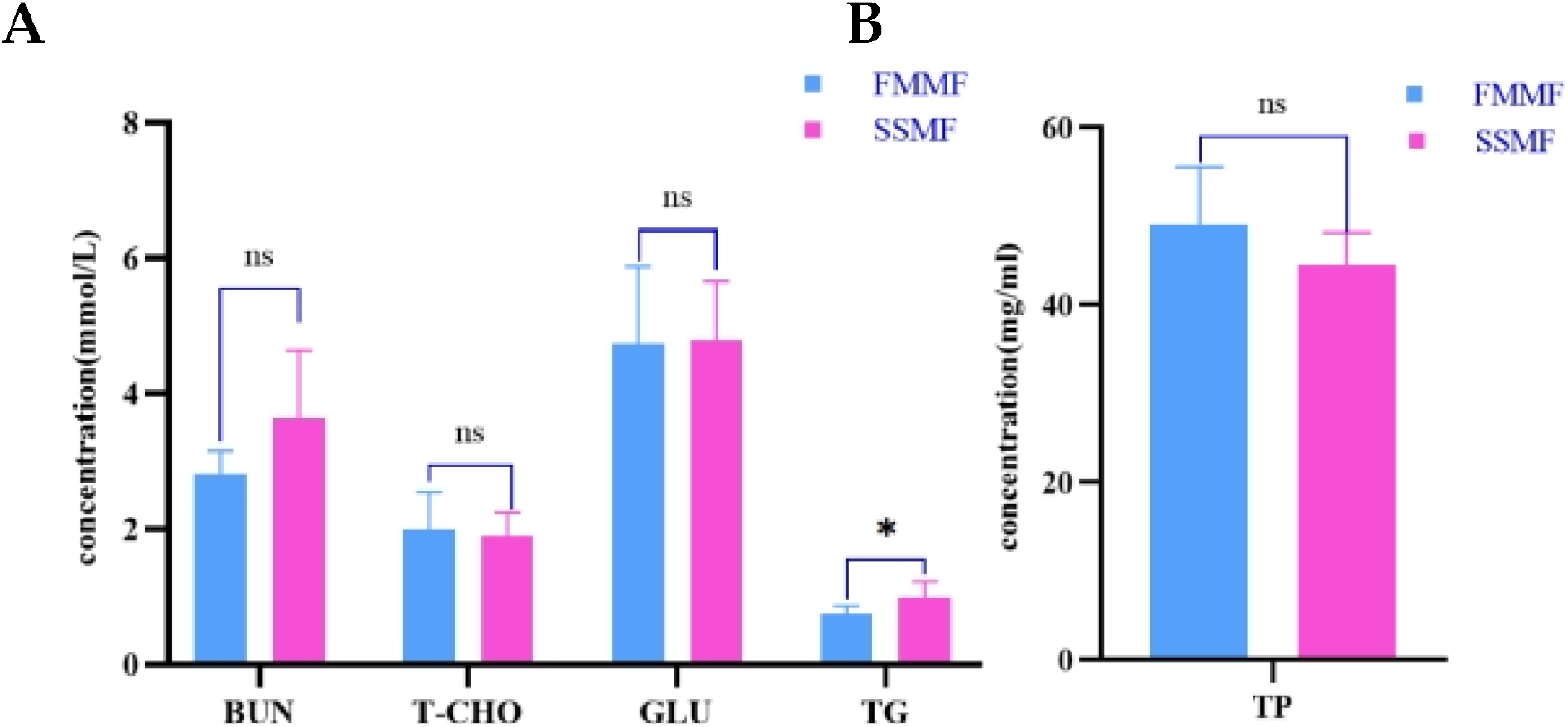
Effects of different feeding methods on serum biochemical indicators of Nanjiang yellow goats 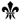 The difference was significant (*P* < 0.05)

### 3.2 Effects of different rearing methods on immunity indexes of Nanjiang yellow goats

As shown in **Fig. 2A**, the serum levels of IgA, IgG, and IgM were higher in the FMMF than in SSMF, but without significant difference. As shown in **Fig. 2B**, the serum levels of IL-2, IL-4, IL-6, and TNF-α were higher in the FMMF than the SSMF, but without significant difference.

**Fig. 2.**
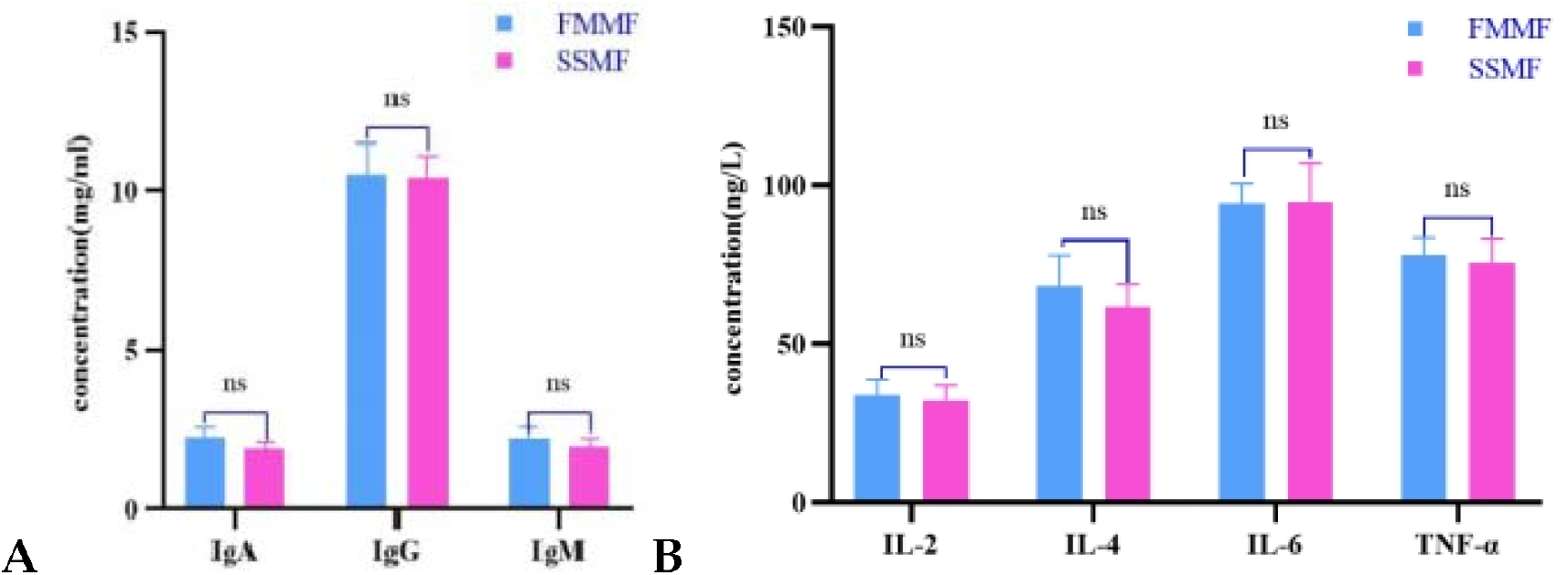
Effects of different feeding methods on serum immune indexes of Nanjiang yellow goats.

### 3.3 Sequencing QC results

After 16S rDNA sequencing, we obtained an average of 85,032 Raw tags for each of the 12 fecal samples of Nanjiang yellow goats, and 72,440 Valid_Tags by quality control, with a quality control efficiency of 85.51%. The statistics of sequencing QC data of each Nanjiang yellow goats sample are shown in the following **Table 1**.

**Table 1.**
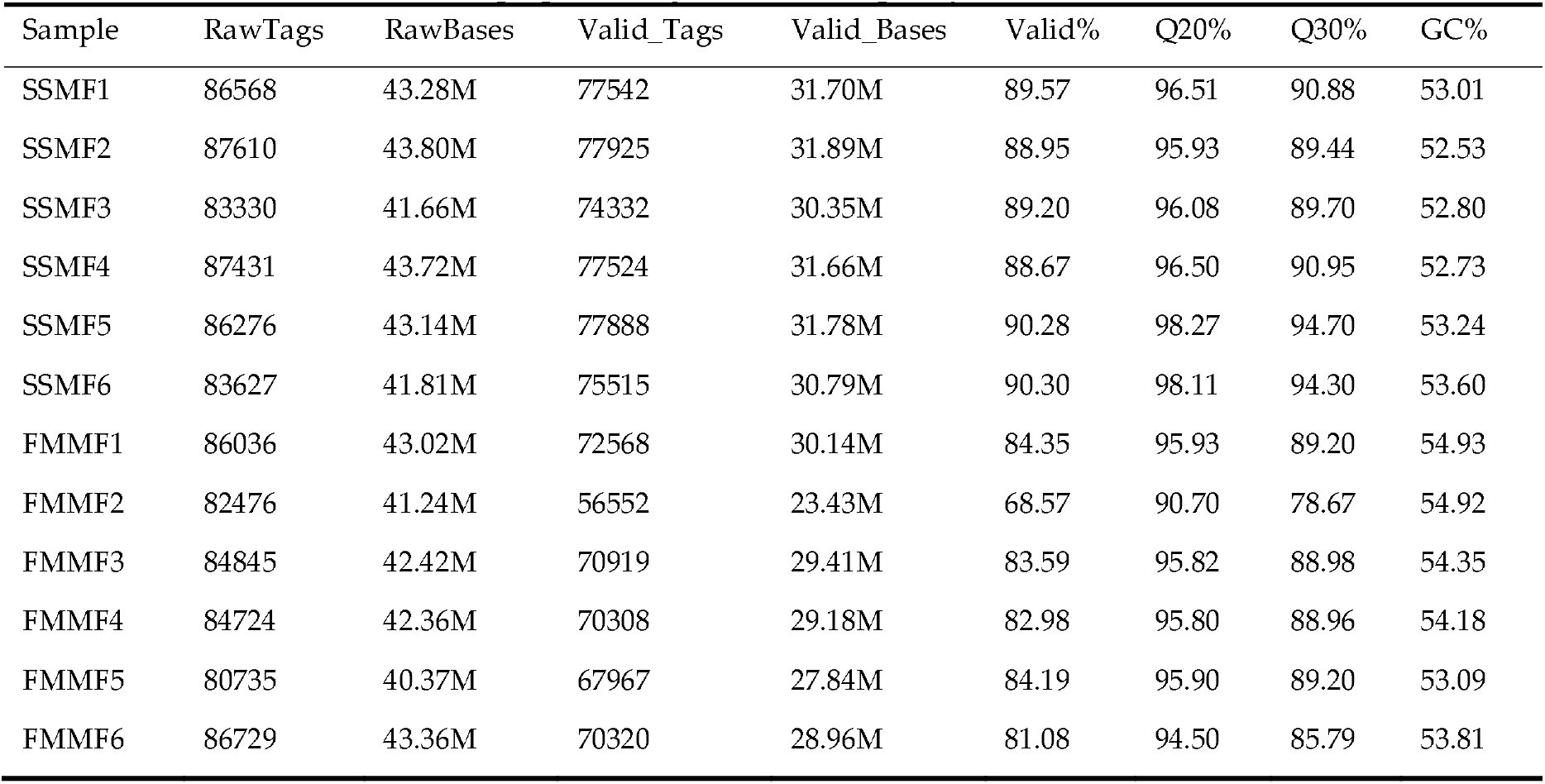
Data preprocessing statistics and quality control.

### 3.4 16S rRNA

#### 3.4.1 Diversity analysis

Universal primers amplified the V3 and V4 regions of the 16S rRNA gene, all the test samples were sequenced and analyzed, and the obtained sequences were optimized and analyzed by OTU clustering according to 97% similarity. This research showed that the relative abundance of information of each sample of the feces of Nanjiang yellow goats in the SSMF group and the FMMF group in different OTUs, and a total of 12 samples clustered into 15,380 OTUs, among which, the 6037 OTUs in the FFMF and 9343 OTUs in the SSMF group. The data showed that the mean values of Chao1 and Shannon’s index (**Fig. 3C**) in the samples from the FFMF of Nanjiang yellow goats were 1055.49 and 8.71, respectively, and the mean values in the samples from the SSMF group were 1737.33 and 9.39, respectively, which showed that the bacterial diversity and the relative abundance in the intestinal tracts of SSMF group were higher than that of the FMMF group, which indicated that the bacteria in the intestine of the SSMF group were more abundant than that of the FMMF group. This shows that the diversity and the relative abundance of bacteria in the intestines of the SSMF of Nanjiang yellow goats were higher than those of the FMMF group of the Nanjiang yellow goats (**Fig. 3D, 3E**).

**Fig. 3.**
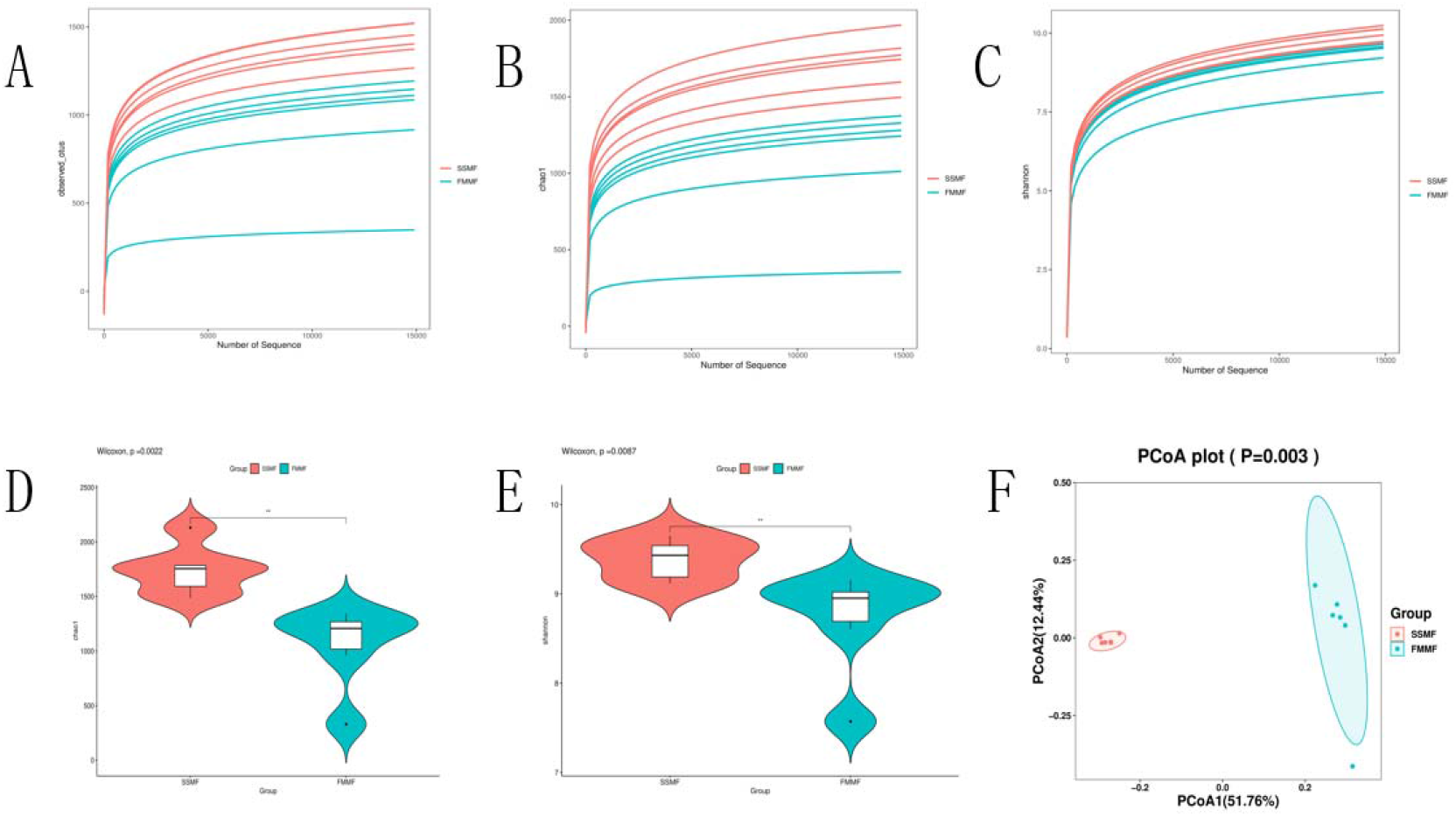
Diversity analysis of intestinal microbiota of Nanjiang yellow goats in SSMF and FMMF Note: A: the average sequencing read length; B: the Chao1 index; C: the Shannon index; DE: the comparative analysis of the Chao1 index and Shannon index of Nanjiang yellow goats in the FMMF and SSMF; F: PCOA analysis.

The average length of the sequenced reads in the SSMF group and the FMMF group was 400-500 kb. From **Fig. 3A**, the dilution curves of the Nanjiang yellow goats samples flattened out after the sequence number reached 15000, which proved that the species in the feces samples of the Nanjiang yellow goats did not increase significantly with the increase in the number of sequenced reads. From the PCOA graph (**Fig. 3F**), we can also find that the intestinal microbiota of 12 Nanjiang yellow goats in the SSMF group and the FMMF group were clustered, which indicated that the intestinal microorganisms in the two feeding methods were obviously different.

#### 3.4.2 Cluster analysis of OUT at different levels

At the phylum level, there were a total of 23 OTUs clustered, with 18 OTUs shared between the SSMF and FMMF groups, accounting for 78.3%. The FMMF group had 3 unique OTUs at the phylum level, comprising 12%, while the FMMF group had 2 unique OTUs, representing 8% (**Fig. 4A**).

**Fig. 4.**
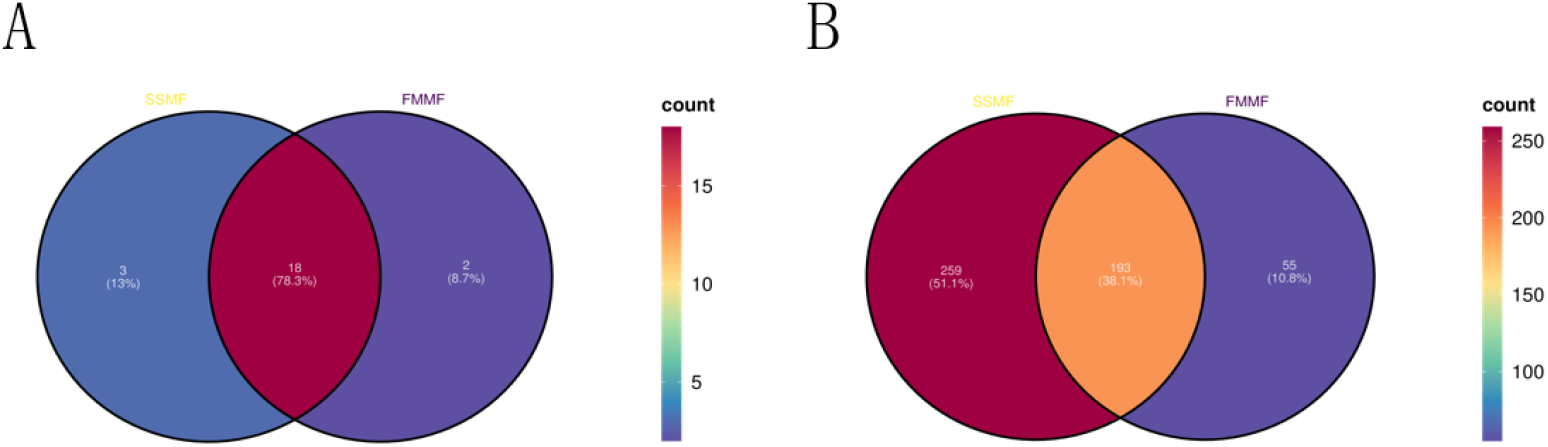
Phylum level vs. genus level OTUs Venn diagrams.

At the genus level, a total of 507 OTUs were co-clustered. Among them, the SSMF group and FMMF group shared 193 OTUs, accounting for 57.3%. The SSMF group had 259 unique OTUs at the genus level, representing 51.5%, while the FMMF group had 55 unique OTUs, comprising 10.8%(**Fig. 4B**).

#### 3.4.3 Analysis of rumen microbiota composition of Nanjiang yellow goats by different feeding methods

We analyzed the microbiota composition in Nanjiang yellow goats at different taxonomic levels, and a total of six major phyla were identified in the feces by 16S rRNA gene sequencing. In the SSMF group, all sequences were categorized into three major phyla (relative abundance of more than 1%) with *Firmicutes* dominating (73.01%, followed by *Bacteroides* (13.83%) and *Verrucomicrobiota* (6.15%). *Firmicutes* were also most abundant in the FMMF group (58.38%) followed by *Bacteroidota* (27.87%), *Patescibacteria* (4.01%), *Verrucomicrobiota* (2.33%), *Proteobacteria* (1.26%). The proportions of *Firmicutes* and *Bacteroides* indicated a prominent escalation in the SSMF group than in the FMMF group, and the proportions of *Spirochetes* and *Proteobacteria* Demonstrated a notable elevation in the FMMF group than in the SSFM group (**Fig. 5A**).

**Fig. 5.**
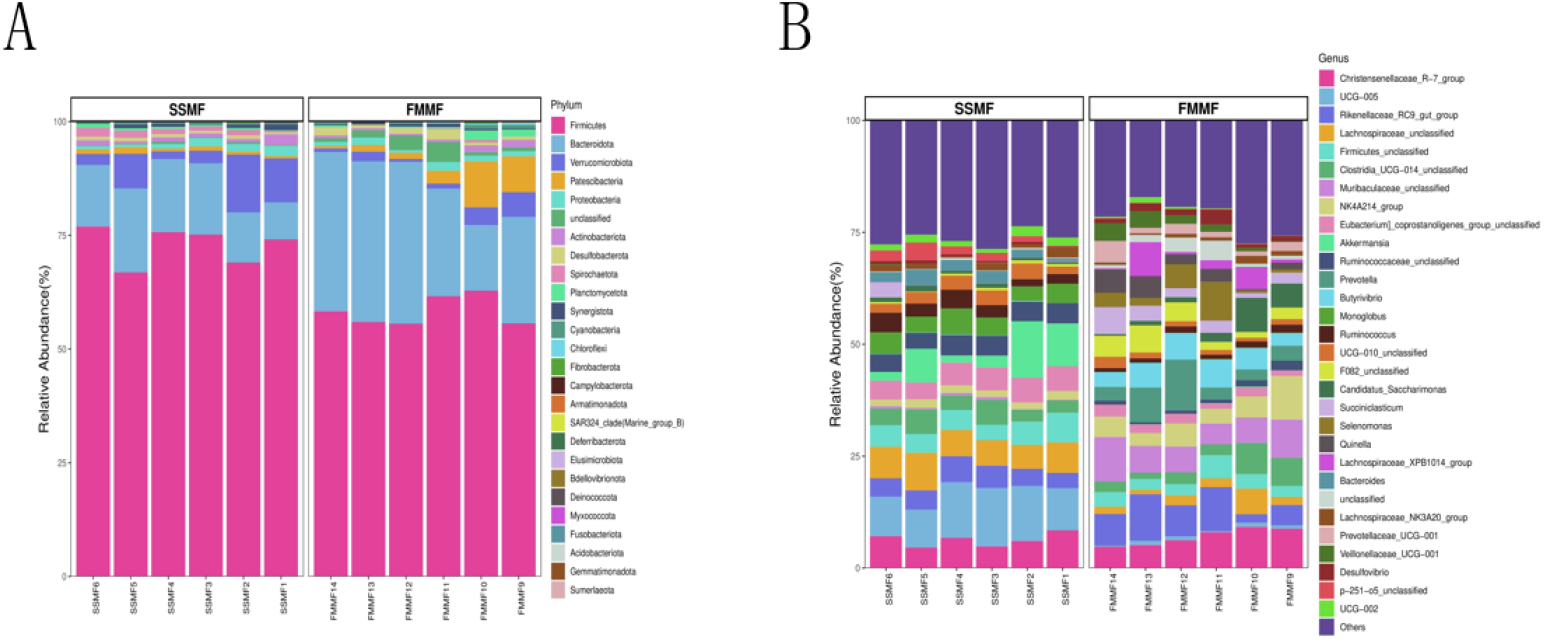
Analysis of rumen bacterial composition of Nanjiang yellow goats in the SSMF group and FMMF group (A: at the phylum level; B: at the genus level)

At the genus level, except for the miscellaneous bacteria (Others), the groups with higher relative abundance in the FMMF group were *Christensenellaceae_R-7_group, Rikenellaceae_RC9_gut_group, Muribaculaceae_unclassified*, and *NK4A214_group*, accounting for 7.0%, 6.78%, 6.74%, and 5.06%, respectively. The relatively higher relative abundance groups in the SSMF group were *UCG-005, Lachnospiraceae_unclassified, Christensenellaceae_R-7_group*, and *Akkermansia*, accounting for 10.77% respectively, 6.53%, 6.30%, and 6.01%, respectively. The relative abundance of *UCG-005, Lachnospiraceae_unclassified, Akkermansia*, and *Firmicutes_unclassifie* was elevated in the SSMF group compared to the FMMF group, while the relative abundance of *Christensenellaceae_R-7_group SSMF* was relatively decreased (**Fig. 5B**).

#### 3.4.4 Differential analysis of the relative abundance of intestinal microbiota in Nanjiang yellow goats by different feeding methods

When performing the analysis of variability in the abundance of microbiota, it was found that a total of 11 species showed prominent differences in relative abundance between the SSMF and FMMF at the phylum level. As shown in **Fig. 6A**, in the FMMF *Armatimonadota, Patescibacteria, Desulfobacterota, Planctomycetota*, and *Chloroflexi* were notably higher (*P*<0.05) and *Fibrobacterotap, Elusimicrobiota, Synergistota, Cyanobacteria* were notably lower (*P*<0.05) compared to the SSMF; and *Spirochaetota, Chloroflexi, Armatimonadota, Deferribacterot* and *Elusimicrobiota, Bdellovibrionota* showed outliers, indicating that compared with the SSMF, the FMMF had a highly remarkable increase in *Armatimonadota*, and *Chloroflexi* (*P*<0.01), a highly remarkable decrease in *Spirochaetota* and *Elusimicrobiota, Deferribacterot* (*P*<0.01).

**Fig. 6.**
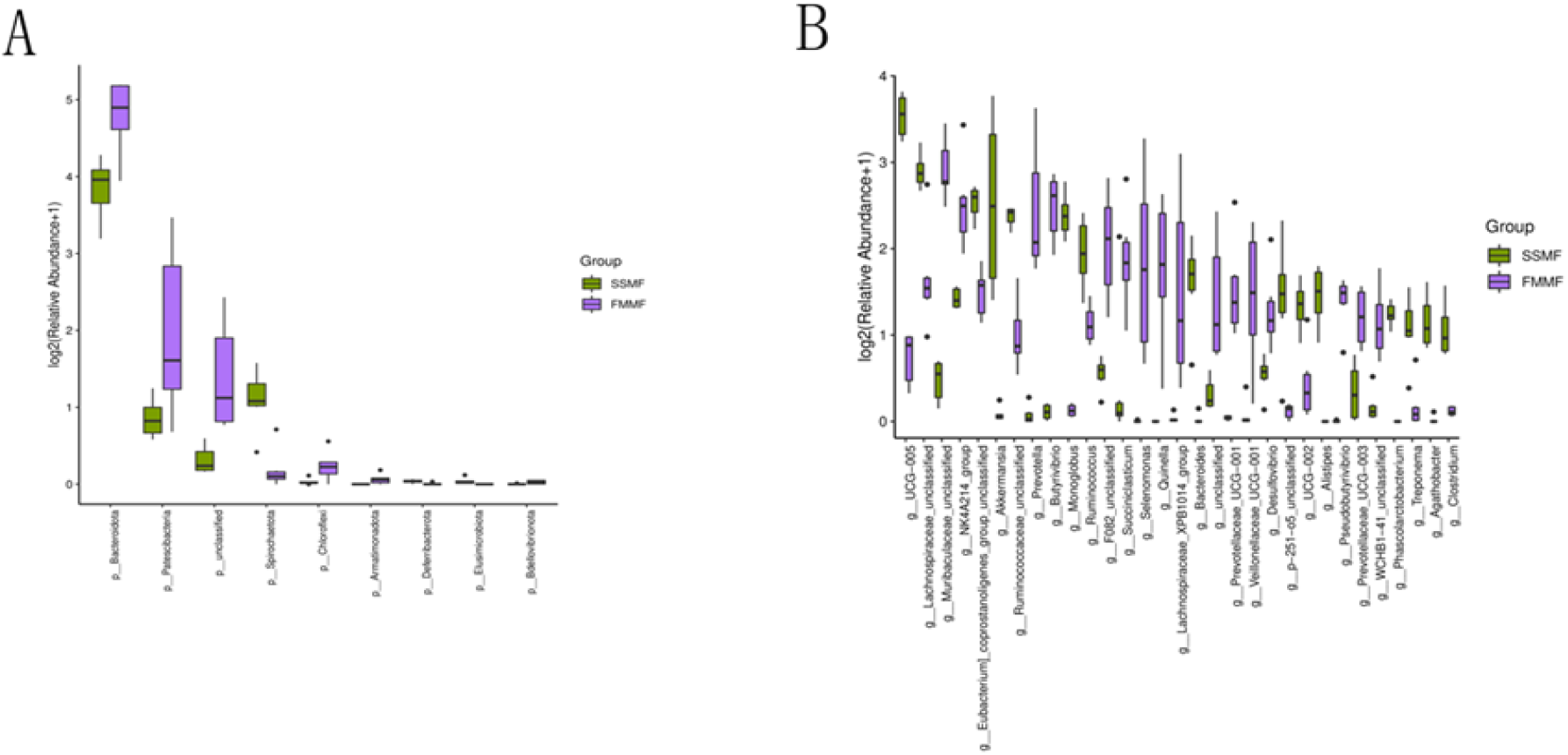
Analysis of the structure of intestinal bacterial composition of Nanjiang yellow goats in the SSMF and FMMF (A: at the phylum level; B: at the genus level)

In the analysis of microbial abundance differences at the genus level, significant differences were observed in the relative abundance of 171 microbial groups between the SSMF and FMMF. As shown in **Fig. 6B**, compared with the SSMF, the relative abundance of *UCG−005, Lachnospiraceae_unclassified, Akkermansia, Monoglobus, Ruminococcus, Bacteroides, UCG−002, Alistipes, Phascolarctobacterium, Treponema, Agathobacter*, and other microbial groups significantly decreased (*P*<0.05) in the FMMF, while the relative abundance of *Muribaculaceae_unclassified, NK4A214_group* and *Prevotella, Butyrivibrio, Succiniclasticum* and *Prevotellaceae_UCG−001* and other microbial groups remarkably increased (*P*<0.05).

### 3.5 Effects of Different feeding methods on intestinal metabolites of Nanjiang Yellow Goats

Intestinal metabolites were evaluated in positive and negative ion modes in Nanjiang yellow goats. A total of 4523 cations were found, of which 1730 were up-regulated and 2793 were down-regulated. A total of 6873 anions were found, of which 2868 were up-regulated and 4005 were down-regulated (**Table 2**). PCA principal component analysis and orthogonal projection latent structure discriminant analysis (OPLS-DA) were performed on all the data from the 12 samples, and as shown in the PCA scatter plot (**Fig.7A**), there was a clear separation of the Nanjiang yellow goats in the FMMF from the SSMF. Further OPLS-DA analysis showed that the FMMF and SSMF groups had great differences in metabolic. (**Fig. 7B**). In addition, the Q2 regression curve had a negative intercept, which indicated the reliability and validity of the OPLS-DA model (**Fig. 7C**). Meanwhile, volcano plot analysis of intestinal metabolites of Nanjiang yellow goats in the FMMF and SSMF revealed notable differences in intestinal metabolites between the two groups (**Fig. 7D**), which indicated that feeding practices could influence the intestinal metabolite composition of Nanjiang yellow goats.

**Table 2.** Number of Upregulated and Downregulated Cations and Anions.

| Ion type | Upregulated | Downregulated | Total |
| --- | --- | --- | --- |
| cationic | 1730 | 2793 | 4523 |
| anionic | 2868 | 4005 | 6873 |

**Fig. 7A:**
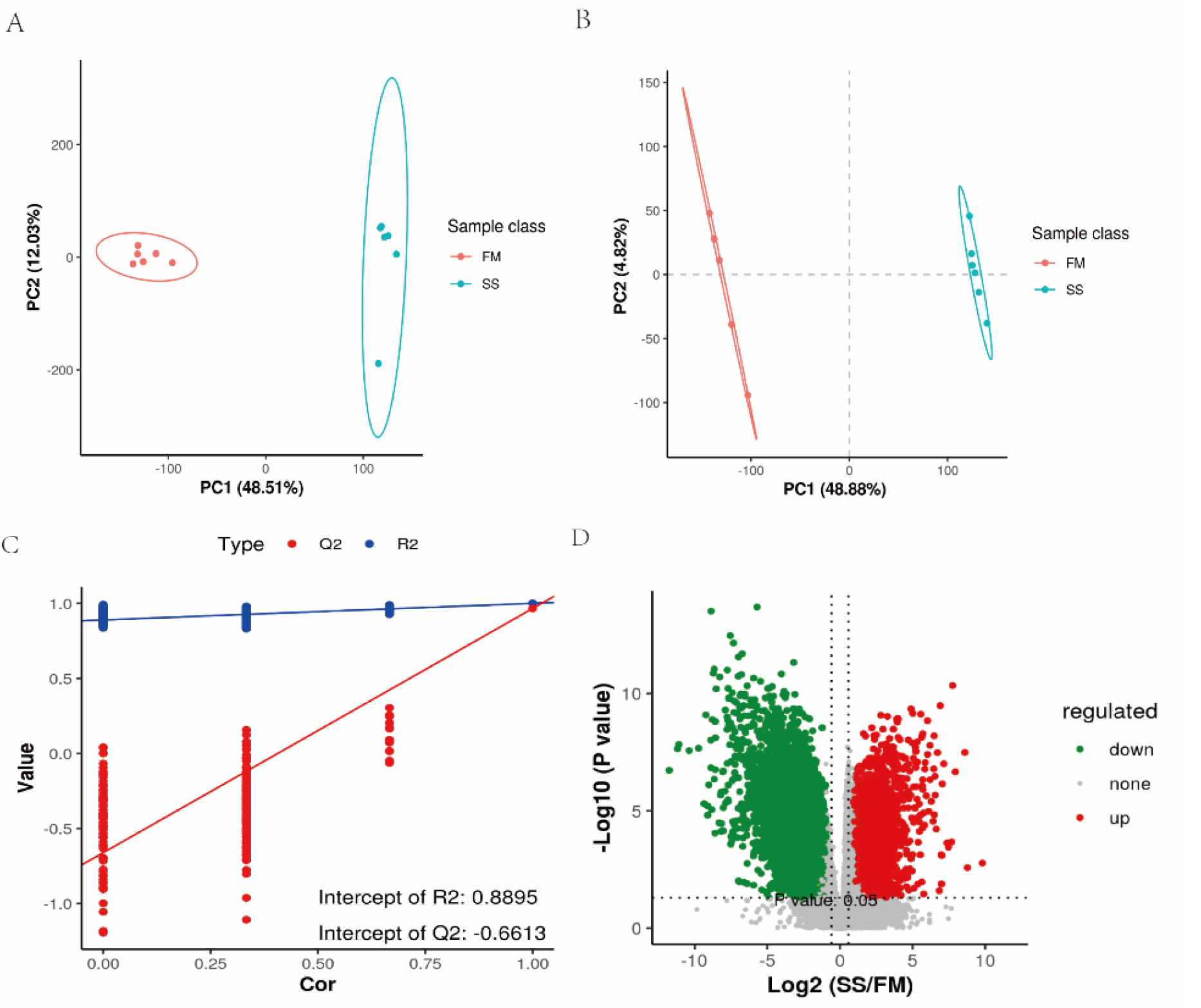
PCA principal component analysis chart; B: Partial Least Squares Discriminant Analysis (PLS-DA) based on intestinal metabolite data; C: PLS-DA validation; D: Differential metabolite volcano plot.

### 3.6 Effects of different feeding methods on intestinal metabolic pathways in Nanjiang yellow goats

Differential metabolites were analyzed by KEGG pathway enrichment to determine whether significant differences occurred in a particular pathway. Differential metabolites were mainly enriched in six categories of pathways, including Cellular Processes, Drug Development, Environmental Information Processing, Human Diseases, Metabolism, Organismal Systems, and Metabolic Pathways. As shown in **Fig. 8**, the most remarkably enriched pathways are Metabolic pathways, including Steroid hormone biosynthesis, Arachidonic acid metabolism, Linoleic acid metabolism, and so on. Eicosanoids and PPAR signaling pathways were also remarkably enriched.

**Fig. 8.**
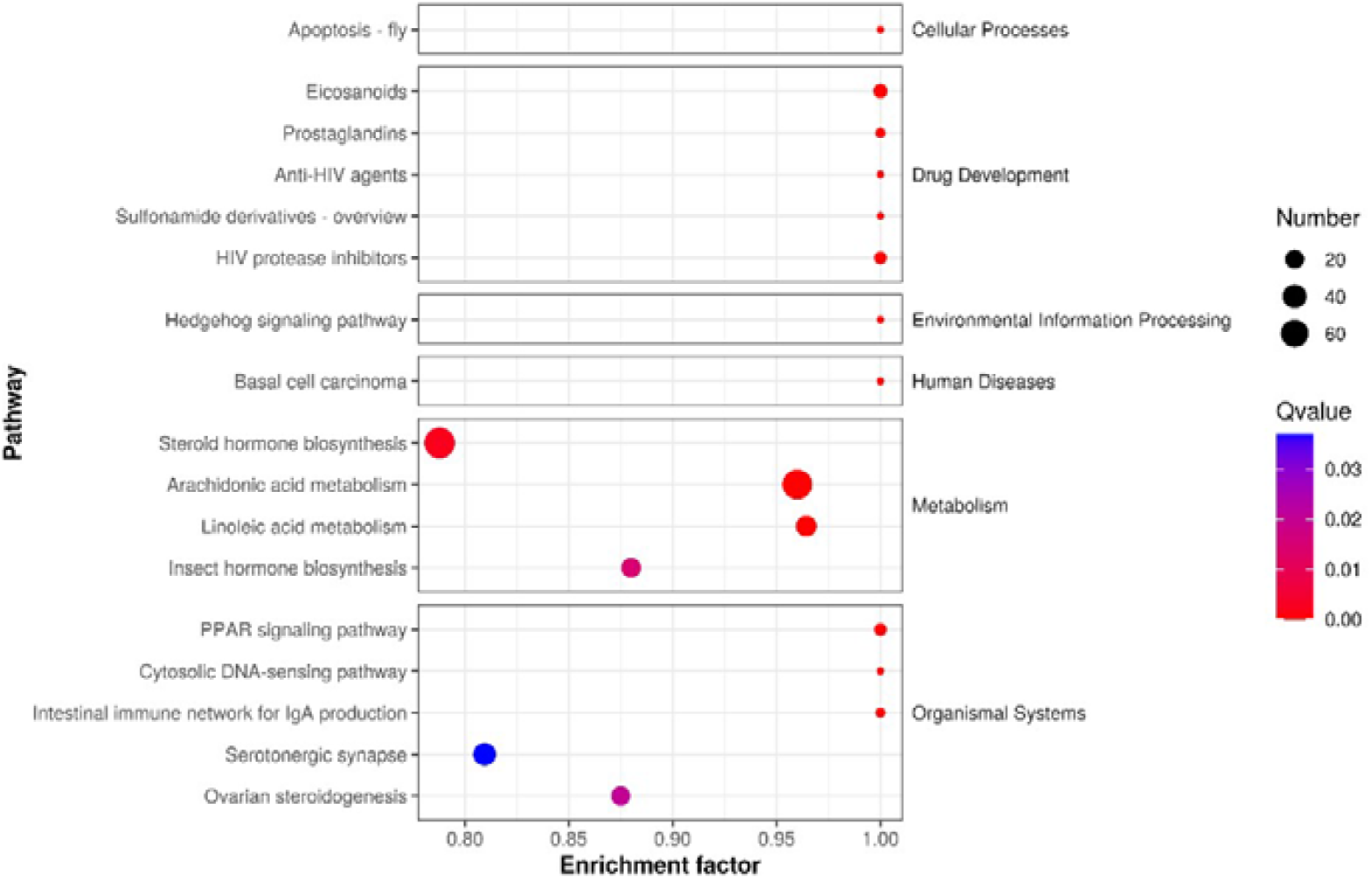
KEGG Pathway Map of Intestinal Metabolites.

### 3.7 Correlation analysis of intestinal microbiota relative abundance and serum immunity indicators

According to the sequencing structure of intestinal microbiota, the top 50 microbial microbiotas with significant differences in relative abundance at the genus level were selected, and after removing unclassified microbial microbiotas, their correlation with serum immune indicators was analyzed.

As shown in **Fig. 9**, the concentration of serum immune factor IL-2 in the serum is showed a strong nagatively correlation with *g__probable_genus_10* and *g__Porphyromonadaceae_unclassified* (*P*<0.05); the concentration of serum immune factor IL-4 is showed a great nagatively correlation with *g__probable_genus_10* and g*__Pseudobutyrivibrio* (*P*<0.05); the concentration of serum immunoglobulin IgA is significant negatively correlated with the relative abundance of *g__Schwartzia* and *g__Faecalibaculum* in the intestines (*P*<0.05), and significantly positively correlated with the relative abundance of *g__Paeniclostridium* in the intestines (*P*<0.05); the concentration of IgM in the serum is showed a strong positively correlation with the relative abundance of *g__Treponema, g__Lachnospiraceae_UCG*-*006, g__p*-*2534*-*18B5_gut_group_unclassified, g__Bilophila, g__Erysipelotrichaceae_UCG* - *003, g__Izemoplasmatales_unclassified* in the intestines (*P*<0.05), and is extremely positively correlated with the relative abundance of *g__Erysipelatoclostridium* in the intestines (*P*<0.01).

**Fig. 9.**
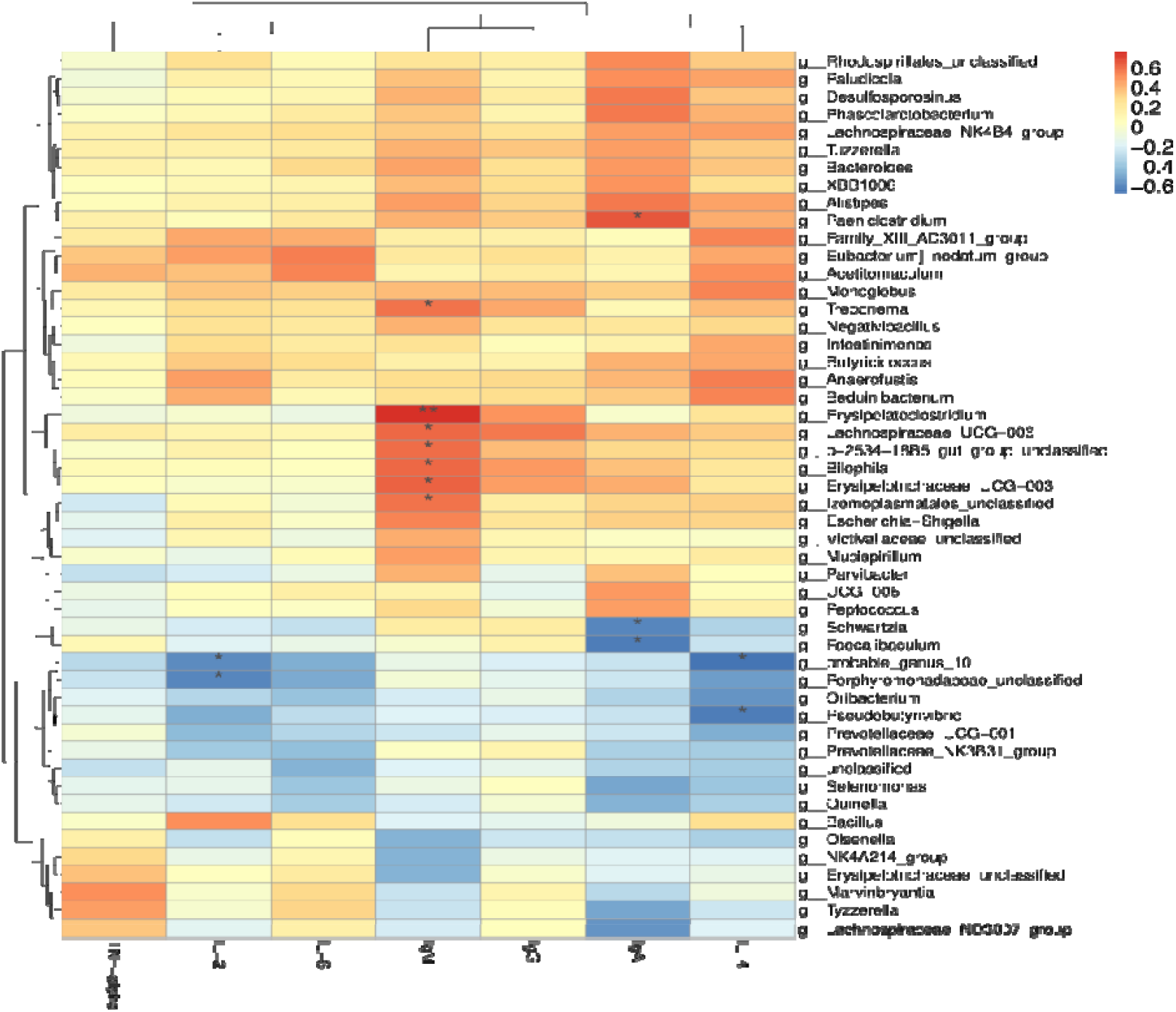
Correlation analysis of serum immunity indexes and intestinal microbiota in Nanjiang yellow goats.

### 3.8 Correlation analysis of gut microorganisms with metabolites

The microbial characteristics and metabolites with remarkably different relative abundances after different feeding methods were identified by multi-omics analysis. Furthermore, we hypothesized that there may be a mechanism linking metabolite concentrations to microbial relative abundance, and this mechanism may be improved by feeding. Therefore, the top 30 differential metabolites were selected for Spearman’s correlation analyses with the top 30 genus-level. Interestingly, most of the differential metabolites were significantly associated with gut microbiota. We visualized all correlations between gut microbial relative abundance and metabolites as plotted in **Fig. 10**. *g__Prevotellaceae_unclassified* with *LysoPG 18:3, 7-Hydroxy-3,4’,8-trimethoxyflavone, Oroselone*, and *FAHFA 22:1* were FAHFA 22:1 were showed a strong positive correlation. ; *g__Bacteroidaceae_unclassified* with *ent-15-Oxo-16 -kauren-19-oic acid, Isolongifolene, 4,5,9,10-dehydro-, 2-methoxy-4-pentadecylbenzoic acid* and *Limaprost* were notably positively correlated; *g__Ruminococcus* was positively correlated with *Azelaic acid, Traumatin, Dodecanedioic acid, PG 30:0* and *Guanine* were had positive correlation. Some significant negative correlations were also found for metabolites and microbiota such as *g__ Desulfobulbus* with *Lupenone, g__ Negativibacillus* with *trans-3-Feruloylcorosolic acid, g__ Bilophila* with (*6Z,9Z,12Z)-6,9,12-Pentadecatrien-2-on*e.

**Fig. 10.**
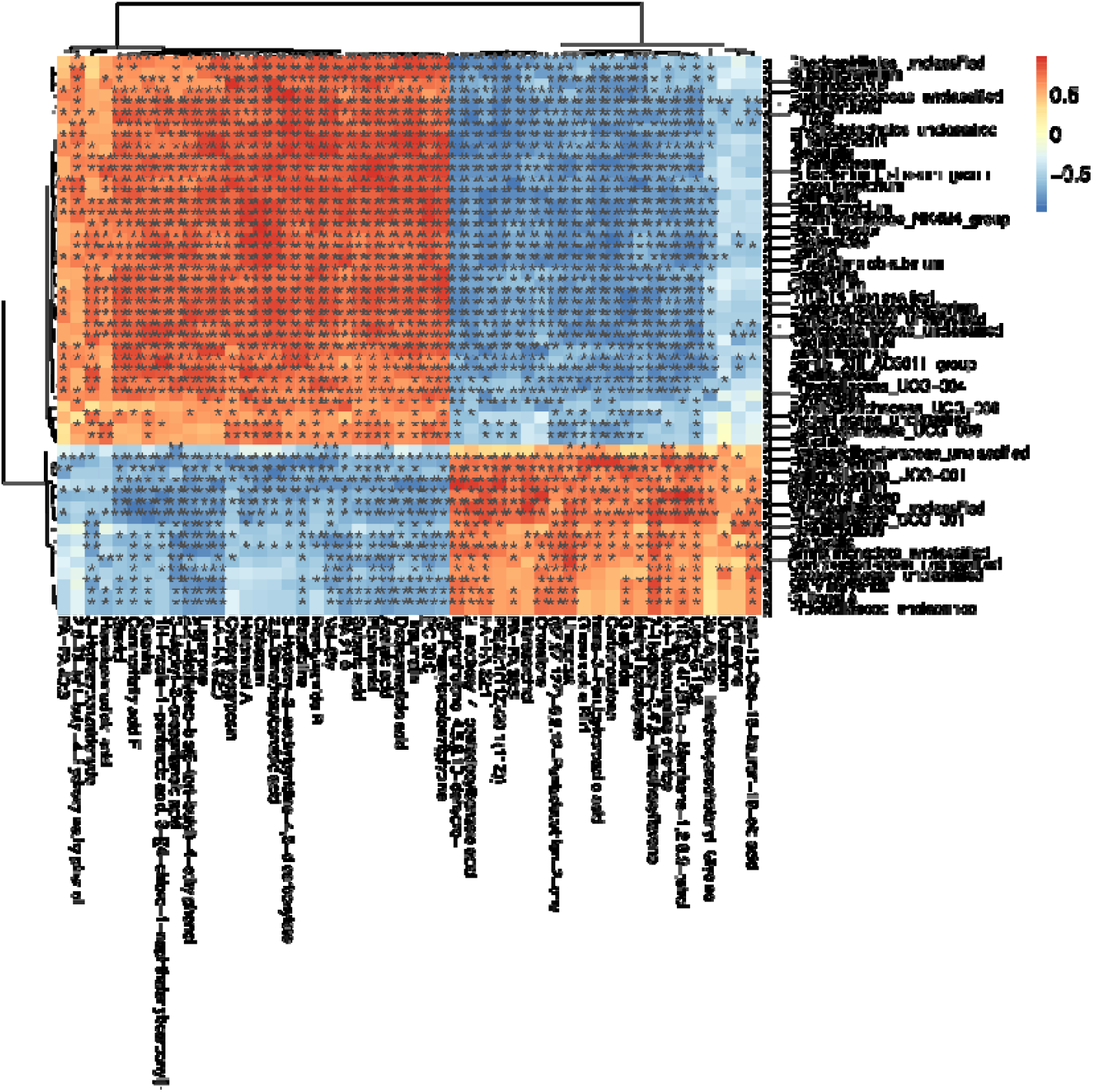
Correlation analysis between intestinal microbiota and metabolites.

**Fig. 11.**
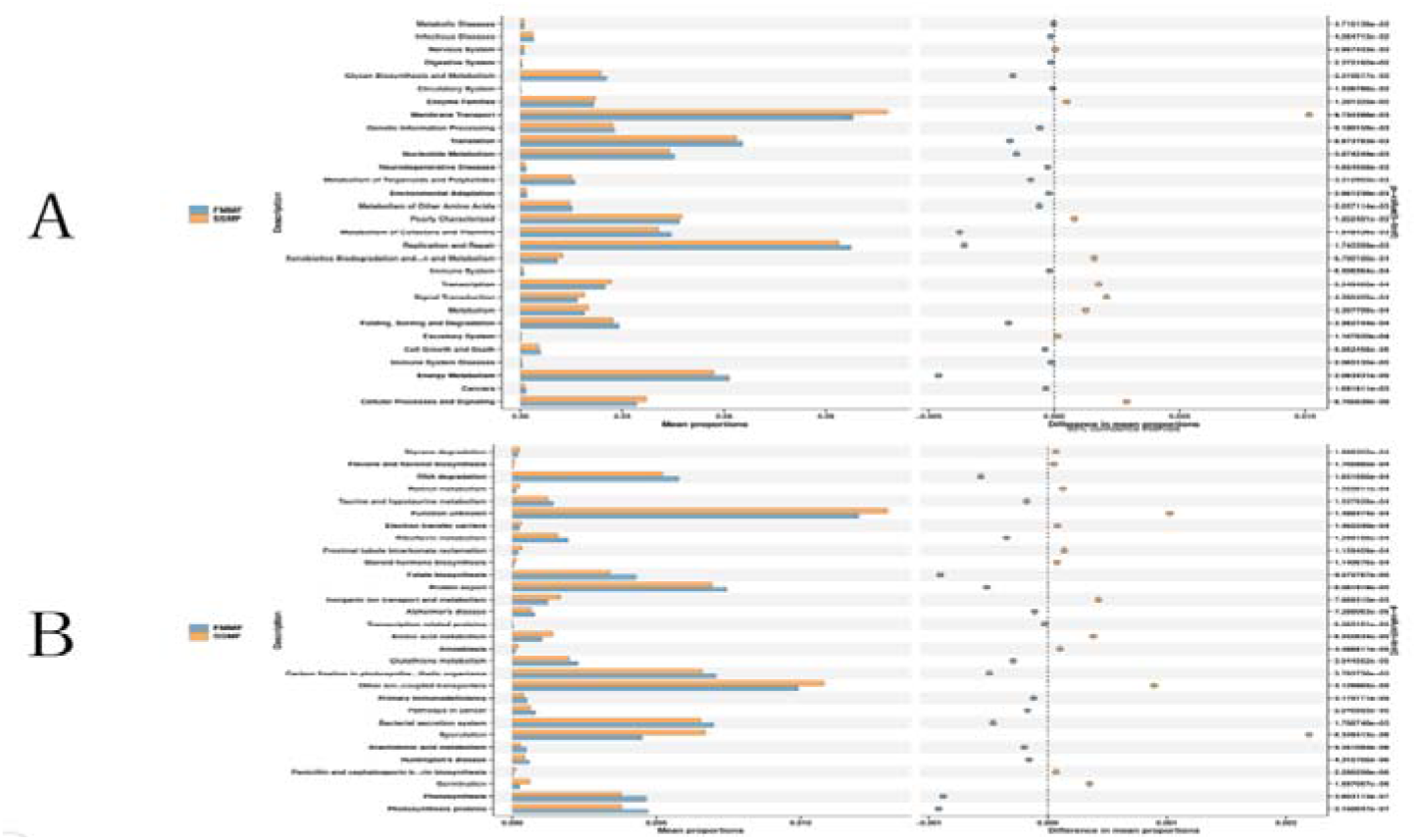
Prediction of metabolic pathways and functions of rumen microbiota (A: KEGG2 level; B: KEGG3 level)

### 3.9 Effects of different feeding methods on the metabolic pathways and functions of the intestinal microbiota of Nanjiang yellow goats

PICUST2 was used to predict the metabolic function of gut microbiota. As shown in (Fig.11A), Membrane Transport was the most abundant metabolic pathway detected at the KEGG2 level, followed by Replication and Repair, Translation, and Energy Metabolism, and Poorly Characterized. The relative abundances of Membrane Transport and Poorly Characterized, Transcription and Cellular Processes and Signaling in the SSFM were significantly higher than those in the FMMF (P < 0.05), while the relative abundances of Translation, and Replication and Repair and Energy Metabolism were significantly lower than those in the FMMF (P < 0.05).

As shown in (Fig.11B), at the KEGG level 3, the most abundant metabolic pathways detected, apart from unknown functions, were Other ion-coupled transporters, Protein export, and Carbon fixation in photosynthetic organisms. Subsequently, Bacterial secretion system, RNA degradation, Photosynthesis proteins, Photosynthesis, and RNA degradation were also identified.

## 4. Discussion

### 4.1. Effect of different feeding methods on serum biochemical indices

Serum biochemical indicators are a method of assessing organ function and disease state of the organism by detecting biochemical substances in the blood. For example, the detection of BUN, creatinine, uric acid, etc. can be used to assess the filtration and excretion functions of the kidneys. Detection of T-CHO, TC, high-density lipoprotein (HDL) cholesterol, low-density lipoprotein (LDL) cholesterol, etc. can be used to assess the metabolism of lipids in the blood and the risk of cardiovascular disease. Changes in serum biochemical markers are usually associated with factors such as environment, age, and different feeding management practices[7]. Serum TG is a fat metabolite that can be broken down by various tissues, and its concentration is closely related to fat metabolism; a high serum TG concentration indicates higher fat utilization and can be regulated by dietary energy levels[8, 9].

In this trial, the levels of TG in the FMMF group were notably decreased compared to those in the SSMF group, which may be related to the diet and amount of exercise in the FMMF group. The FMMF group generally consumed grass as their staple food, which is high in carbohydrates and low in fats, which resulted in lower TG levels in the FMMF group. Further, the FMMF group is going to exercise more than the SSMF group, which depletes the body of energy, including fat, resulting in lower TG in the FMMF group. TG in adipose tissue releases several important immunomodulatory factors, and low levels may lead to decreased immune function. This may make goats more susceptible to infection and disease. TG can also affect the synthesis and balance of sex hormones, and low levels may have some effect on reproductive function in goats and this may lead to reduced fertility or reproductive problems.

Relevant studies have shown that different feeding methods have a certain impact on protein metabolism while feeding energy levels by affecting the metabolism and thus affecting the concentration of GLU in the serum when the energy intake of the animal body is insufficient, the concentration of GLU in the serum will drop significantly, thus reflecting the decline in the utilization of feed nutrients by the animal[10]; Zhang et al. found that serum GLU levels in overgrazed goats were significantly lower than those in the lightly grazed group[11], this is consistent with that we found the serum GLU content is more in SSMF group. And Grigol et al. in comparing the effects of different grazing times on the health status of dairy cows, found that an increase in the time spent on grazing led to a decrease in the BUN levels of the animals[12]. This result is also consistent with the outcome of this research that the serum level of BUN in FMMF was lower than in the SSMF group. Since the results showed that there was no significant difference in BUN level and GLU level between FMMF and SSMF, it can be deduced that barn-feeding and grazing had no or little effect on the body of Nanjiang yellow goats on fat utilization, energy intake, and metabolism. GLU is an important source of energy, which is mainly produced through the metabolism of carbohydrates. Overgrazing may lead to insufficient carbohydrate content in livestock feed, thus affecting the availability of GLU. And lack of GLU may lead to energy deficiency in livestock, affecting growth and performance. BUN is one of the indicators to assess renal filtration and excretion function. Abnormalities in BUN levels may indicate abnormalities in the nutritional status and renal function of livestock.

We found that the T-CHO content and TP content of the FMMF group were higher than those of the SSMF group in this research. Serum T-CHO can reflect the nutritional value of feed, animal physiology, growth, and development level [13], cholesterol is a kind of lipids, mainly synthesized by the animal body. The T-CHO content of FMMF in this experiment was lower than SSMF, and it can be inferred that the utilization rate of feed nutrients, the growth development level of grazing goats are relatively lower than that of barn-feeding goats. The TP content in serum reflects the absorption, synthesis, and catabolism of proteins, as well as the ability of the liver to synthesize proteins. The lack of significant difference in TP between SSMF and FMMF groups in this paper may result from the fact that there is no great difference in their nutrient intake and efficiency of digestion and absorption.

### 4.2. Effect of different feeding methods on serum immunity indexes

Ig is a class of protein molecules found in the body and an important part of the body’s immune system. Ig are produced by immune cells (B-lymphocytes) and function as antibodies in the body. Among them, IgA mainly plays a role in protecting mucosal surfaces from pathogens. IgG has a wide range of antibody activity that neutralizes pathogens and promotes inflammatory responses, etc. IgM has a strong ability to neutralize pathogens.IL-2, IL-4, IL-6, and TNF-alpha are cytokines in the immune system, and they play an important role in regulating and modulating immune responses. IL-4 promotes the B-cells to differentiate into antibody-producing plasma cells and enhances the production of immunoglobulin E (IgE). TNF-α is produced by a variety of immune cells and is an important inflammatory mediator that plays a key role in inflammatory responses and immune regulation.

Environmental factors and dietary structure can influence the immune structure of the body as well as cell maturation and function[14]. A study on the effect of housing and semi-housing on serum biochemical indexes in Jersy cattle found that the serum IgM content of the semi-housing group was significantly higher than that of the housing group[15], a result that is consistent with the present study in which the serum IgM content of the FMMF group was higher than that of the SSMF group. Our results indicate that the IgA, IgM, and IgG contents of the FMMF group were higher than those of the SSMF group, and Li Qin et al. showed that the amount of exercise is the main factor affecting blood immunity indexes[16]. According to this paper, the FMMF group could move freely and exercise, which helped to improve the function of the circulatory system and the activity of immune cells. Exercise promotes lymphatic fluid circulation, which enhances the transportation of immune cells and antibody production. Secondly, it may also be related to the feeding environment. The FMMF group lives in a natural environment and is exposed to more microorganisms and environmental stimulants. These stimulants can promote the activation of the immune system and antibody production[17], thus increasing IgA, IgM, and IgG levels in goats. It has been shown that serum interleukin 4 (IL-4) levels are significantly decreased in opened house feeding lambs compared to closed house feeding lambs[18], which is consistent with the results of the present study that IL-4 levels were higher in the FMMF group than in the SSFM group.

The levels of TNF-α are related to several factors. First, TNF-α is closely interrelated to an inflammatory response. When the body suffers from infection, injury, or other inflammatory stimuli, immune cells release TNF-α, which promotes the occurrence and development of inflammatory responses[19]. From the validation of these results, the TNF-α content in the serum of the FMMF group was higher than that of the SSMF group, which may be due to the fact that the harmful gases in the grazing air of the Nanjiang yellow goats were higher than that within the barn-feeding animals, and the long-term inhalation of harmful gases by the goats led to inflammation occurring in the body, which in turn caused the elevation of the TNF-α content in the serum.

### 4.3. Effect of different feeding practices on gut microorganisms

The majority of the mammalian gut microbiota is acquired from the environment from the time it is born, and its composition is largely determined by factors such as its species, age, feeding practices, environment, and disease. Most researchers would argue that host species play a greater role than environmental factors in shaping the gut microbiota, especially when there are large taxonomic differences between host species. To date, the rumen has been the focus of most researchers because it is the primary site of feed fermentation. The gut microbiota of ruminants has been less studied. However, it has been shown that energetically important microbial products, including VFA, can be produced in the gut of ruminants[20, 21]. Bacteria occupy a major portion of the intestinal microbiota and their presence is essential for the health of dairy cows. They help in the fermentation and degradation of plant polymers by secreting various enzymes which help in the fermentation and degradation of plant polymers[22, 23]. The identification of these bacteria and their unique functions have been the focus of many studies. With advances in next-generation sequencing, microbiotechnology, culture-free methods, and genetic engineering, it has become easier to study the role of commensal microbiota in host metabolism. The predominant colony in the gut is *Firmicutes*, followed by *Bacteroidota, Patescibacteria*, and *Proteobacteria*.

We usually use the feces and rumen contents of ruminants to study the microbiome of their gastrointestinal tract[24, 25]. In this experiment, the feces of Nanjiang yellow goats were used to characterize their intestinal microbiome. Our results showed that in the SSMF, *Firmicutes* was the highest relative abundance followed by *Bacteroides* and *Verrucomicrobiota*. In the FMMF, *Firmicutes* were also the most abundant, followed by *Bacteroidota, Patescibacteria, Verrucomicrobiota*, and *Proteobacteria*. The proportion of *Firmicutes* and *Bacteroides* was significantly higher in the FMMF than in the SSMF, and the proportion of *Spirochetes* and *Proteobacteria* was significantly higher in the grazed group than in the SSMF. Ley et al. investigated the evolution of mammalian gut microbes and said that the dominant phylum was *Firmicutes* followed by the *Anaplasma* phylum and *Aspergillus* [26, 27].

### 4.4. Effect of different feeding methods on gut metabolites

Metabolites are endogenous small molecular compounds, such as amino acids, sugars, lipids, nucleic acids, etc., that participate in the metabolism of an organism and maintain its normal growth and developmental functions, and the production and accumulation of these metabolites reflect the state of the biological system and the dynamic process of the whole system[26, 28]. Metabolomics technology can use different analytical methods, such as mass spectrometry, nuclear magnetic resonance (NMR), and liquid chromatography, to detect and quantify metabolites, thus revealing the metabolic state and function of an organism. Metabolomics has been widely used in many research areas such as nutritional assessment of livestock and poultry, environmental stress, livestock product safety diagnosis, growth and development, and meat quality assessment.

In this research, we assessed the effects of different feeding practices on the intestinal metabolome using an untargeted metabolomics technique, i.e., LC-MS. Metabolites drive key cellular functions, such as energy production and storage, which can further affect intestinal developmental status. Therefore, it is important to explore the effects of different feeding practices on intestinal metabolites to reveal the intestinal development mechanism of Nanjiang yellow goats. Wang et al. found that the levels of 43 of these serum metabolites were notably elevated in high-growth goats by comparing the serum metabolism differences between high-growth and low-growth goats, including D-ornithine, l-glutamine, L-histidine, myosin, lysophosphatidylinositolase, DCTP, and hydroxylysine; the levels of 82 serum metabolites were remarkably higher in low-growth rate goats, including p-salicylic acid and deoxycholic acid 3-glucuronide[28]. Wang et al. explored the effects of dietary energy and protein on the rumen bacterial composition and rumen metabolites of the velvet goats and found that there was a significant difference in the metabolites between the high-energy, high-protein group and the control group. The main differential metabolites were amino acids, peptides, and analogs[29]. The characteristics of the combination of microbiome and metabolome in cows with different intake levels were investigated, and it was found that the stearic acid content of cows in the high-intake group was significantly higher than that in the low-intake group[30].

Based on this research, a total of 4523 cations were found, of which 1730 were up-regulated and 2793 were down-regulated, and a total of 6873 anions were found, of which 2868 were up-regulated and 4005 were down-regulated. These mainly included Prostaglandin G1, Dexamethasone, LysoPE 18:1, and Heptadecanoic acid, suggesting that different feeding practices may result in outstanding differences in gut metabolite levels. Heptadecanoic acid has been found to account for a small fraction of total saturated fatty acids in milk and ruminant meat and is recognized as a milk fat biomarker as well as a possible lamb flavor substitute [31, 32]. Therefore, in this report, it was hypothesized that feeding practices altered the flavor of Nanjiang yellow goats by affecting the content of Heptadecanoic acid. Further by KEGG analysis, we found that the most notably enriched surrogate pathways included Steroid hormone biosynthesis, Arachidonic acid metabolism, and Linoleic acid metabolism.

By analyzing serum reproductive hormones and ovarian genes of pubertal female goats, we found significant enrichment including estrogen signaling pathway, steroid hormone biosynthesis, and cAMP signaling pathway[33]. Investigating the differences in shaping the gut microbiota and regulating metabolism in cow’s milk-based infant formulas, goats’ milk-based infant formulas, and mixed-milk-based infant formulas compared to pasteurized breast milk, their metabolomics identification showed that goats’ milk-based infant formulas mainly emphasized bile acid biosynthesis, arachidonic acid metabolism, and steroid biosynthesis metabolic pathways[34]. To explore the biological activity of endogenous melatonin, melatonin-rich dairy goats overexpressing acetylserotonin-O-methyltransferase (ASMT) were successfully prepared using pBC1-ASMT expression vector and microinjection of prokaryotic embryos, and metabolic analyses revealed that the transgenic goats had a unique arachidonic acid metabolic pattern[35].

### 4.5. Correlation analysis of different feeding methods on the relative abundance of intestinal microbiota and serum immunity indexes in Nanjiang yellow goats

The intestinal tract is the largest digestive organ in the human body. It is colonized by, and consistently exposed to, a myriad of microorganisms, including *bifidobacteria, lactobacillus, Escherichia coli, enterococcus, clostridium perfringens*, and *pseudomonas* [36]. There are constant interactions between the intestinal microbiota and the immune system. In addition, the intestinal microenvironment can be reconstructed by using probiotics or microbiota transplantation. Studies showed that the adhesion of bacteria to the gut epithelium is believed to help establish colonization in the digestive tract [37, 38]. In the early stages of life, proper intestinal colonization of specific microbiota can stimulate the maturation of intestinal mucosa-associated lymphoid tissue [39]. The intestinal mucosal immune barrier is composed of humoral immunity and cellular immunity. The immune cells and cytokines in the intestinal mucosa maintain intestinal homeostasis by participating in innate immunity and adaptive immunity [39].

This research found that the relative abundance of *g__probable_genus_10, g__Porphyromonadaceae_unclassified, g__Pseudobutyrivibrio* in the intestine was negatively correlated with the concentration of serum immune factors IL-2 and IL-4, indicating that these intestinal bacteria may inhibit the production of plasma cells and affect the production of IgM. As a kind of intestinal microorganism, *g__Pseudobutyrivibrio* may affect the host’s immune system through its metabolites or interaction with intestinal epithelial cells. This effect may include reducing the ability of immune cells to produce cytokines such as IL-2 and IL-4, thereby affecting the balance and function of the immune system. In this paper, there was a remarkable positive correlation between the relative abundance of *g__Erysipelatoclostridium* and the concentration of IgM in serum. There is also some research showing that the *Erysipelatoclostridium ramosum* is an intestinal commensal bacterium, some of which can produce IgA1 and IgA2 proteases [40]. Therefore, these gut bacteria may regulate the response and function of the immune system by affecting the structure and function of the gut microbiota.

The results of this research showed that among the top 50 microbiota with outstanding differences in relative abundance, there were 8 microbiota with significant positive correlation with IL-2, IL-4, IgM, and IgA, and 5 microbiota with significant negative correlation. The effects of different feeding methods on serum immune indexes and intestinal microbial relative abundance of Nanjiang yellow goats were also quite different. Based on the characteristics of blood immune indexes and intestinal microbiota, it was found that grazing feeding may increase the relative abundance of intestinal beneficial microbiota and increase the concentration of immune protein IgM, IgA, and immune factors IL-2 and IL-4 in serum, thereby enhancing the immune performance of Nanjiang yellow goats.

### 4.6. Correlation analysis of gut microbes with metabolome

To date, multi-omics analysis has been widely applied in various animals such as pigs, cattle, sheep, and poultry. Its primary applications include the assessment of animal nutritional levels, growth and development, reproductive performance, and meat quality evaluation [41-45]. Heinritz, based on microbiome and metabolome findings, discovered that pigs fed a low-fat/high-fiber diet (LF) had higher levels of *lactobacilli, bifidobacteria* (*P*<0.001), and *Faecalibacterium prausnitzii* (*P*<0.05) compared to those on a high-fat/low-fiber (HF) diet. In the HF diet, there were more proteins related to *Enterobacteriaceae*, and proteins for polysaccharide decomposition were almost entirely derived from *Prevotellacea* [46]. **Xue** found that Prevotella and Succinimonas amylolytic were enriched more than 6 times in the rumen of dairy cows with high milk yield and milk protein content by correlation analysis of rumen microorganisms and metabolites in different dairy cows. As succinate-producing bacteria in the rumen of cattle, they were positively correlated with VFAs concentration, indicating their important role in VFAs biosynthesis[47]. **Shen** and colleagues investigated changes in the lung microbiota and metabolome of broiler chickens exposed to particulate matter (PM) collected from chicken coops. They observed notable changes in *α* and *β* diversity induced by PM. Using untargeted metabolomics, they identified several differential metabolites induced by PM and related to different bacteria. They found that disturbances in the microbiome and metabolome might play a crucial role in the mechanism of PM-induced lung damage in broilers [48].

We explored the impact of different feeding methods on the gut microbiota and metabolites of Nanjiang Yellow goats using metabolomics and microbiomics, and further analyzed the correlation between gut microbiota and metabolites using the Spearman analysis method. Can, in a goats model fed a high-concentration diet, found that the strongest positive correlation in bacteria and metabolites was between hexadecanoic acid and Bulleidia, and the strongest negative correlation was between L-tyrosine and Oscillospira [49]. Huang and others, through microbiome and metabolomics studies, revealed the impact of different feeding systems on the growth and rumen development of yaks and performed correlation analysis of gut microbiota and metabolites. They found that the dynamic fluctuations of some metabolites were closely related to the relative abundance of various microbial communities. Among them, the genus *Ruminococcaceae_UCG 001* was negatively correlated with dopaquinone, adenine, and guanosine but positively with xanthine, L-tyrosine, and hydroxyphenyllactic acid. The genus *Rikenellaceae RC9* gut group was positively correlated with dopaquinone, adenine, guanosine, and inosine, but negatively with xanthine, L-tyrosine, acetylcholine, and hydroxyphenyllactic acid [50]. This research found that *g__Prevotellaceae_unclassified* was outstandingly positively correlated with LysoPG 18:3, 7*−*Hydroxy*−*3,4’,8*−*trimethoxyflavone, Oroselone, and FAHFA 22:1. *g__Prevotellaceae_unclassified*, primarily present in adult animals, plays a crucial role in the degradation of fiber, starch, or protein, and in ruminants, mainly breaks down plant fibers. LysoPG 18:3, through emulsification and its vital physiological functions, promotes the digestion and absorption of fats, proteins, and other nutrients, reduces diarrhea, improves feed utilization efficiency, and promotes the growth of young animals. The term “FAHFAs” was introduced for bioactive lipids discovered in fat cells with potential anti-diabetic and anti-inflammatory properties. FAHFAs are a large class of endogenous bioactive lipids, now presenting new opportunities for treating diabetes and inflammatory diseases [51, 52]. *g__Ruminococcus* was outstandingly positively correlated with Azelaic acid, Traumatin, Dodecanedioic acid, PG 30:0, and Guanine. *g__Ruminococcus* comprises two strong fiber-digesting bacteria, *Ruminococcus albus* and *Ruminococcus flavefaciens*, capable of producing a large amount of cellulase and hemicellulase. The highe relativer abundance of rumen coccus corresponds to a higher fiber content, likely due to substrate-induced effects. Guanine is crucial in the field of ruminant nutrition, particularly for quantifying microbial protein synthesis and degradation in the rumen, as well as for all protein evaluation systems. Guanine is used as an internal microbial marker for determining microbial protein synthesis [53, 54].

Based on the above data, we can speculate that different feeding methods may have a significant impact on the gut microbiota and metabolic products of Nanjiang Yellow goats. This finding highlights the crucial role of different feeding methods in shaping the gut microbial composition, thereby affecting the types and concentrations of metabolites. Understanding how to adjust feeding methods to improve or maintain the health of livestock holds significant importance.

### 4.7 Prediction of metabolic pathways and functions of gut microbiota by different feeding practices

The complex relationship between gut microbes and metabolic function has been the focus of numerous studies, and the metabolic activity of gut microbes is critical for maintaining host homeostasis and health. Gut microbes play a key role in nutrient metabolism, foreign body and drug metabolism, and maintenance of the gut barrier. The important contribution of the gut microbiota to human metabolism, which provides enzymes not encoded by the human genome, has also been emphasized[55, 56]

From this experiment we conclude that Membrane Transport was identified as the most abundant metabolic pathway at KEGG2 levels, followed by Replication and Repair and Translation and Energy Metabolism. Notably, the relative abundance of Membrane Transport, Cellular Processes and SignalingTranscription was remarkably higher in the SSMF than in the FMMF. In contrast, Translation, Replication and Repair, and Energy Metabolism were remarkably lower in the FMMF [57].

At the KEGG level 3, data also revealed notable differences in certain metabolic pathways between the SSMF and FMMF. For instance, the relative abundance of Other ion-coupled transporters and Sporulation was significantly higher in the SSMF compared to the FMMF. This might suggest that microbes in the barn-feeding environment rely more on ion-coupled transport mechanisms for nutrient acquisition, while microbes in the grazing environment may have more opportunities for sporulation to cope with environmental instability [58].

On the other hand, in the FMMF, the relative abundance of Bacterial secretion system, Carbon fixation in photosynthetic organisms, Protein export, Photosynthesis proteins, and Photosynthesis was remarkably lower than in the SSMF. This could reflect a lower dependence of microbial communities in the grazing environment on photosynthesis and protein secretion, aligning with the potential prevalence of nutrition from plant sources in the grazing environment [59].

In summary, the metabolic functions of gut microbiota under different feeding conditions provide intriguing insights into how microbial communities adapt and diversify their metabolism in response to dietary and environmental changes. Further research could delve into the mechanistic basis of these metabolic shifts and their specific implications for host health and nutrition in diverse environments and under different feeding methods[60].

## 5. Conclusion

Through comparative analysis of gut microbiota structure and serum immune indicators in Nanjiang yellow goats under different feeding methods, differences were observed between the SSMF and the FMMF in terms of serum biochemical indicators, immune indicators, and gut microbiota and metabolic indicators. There was a significant impact on the serum TG levels. Compared to FMMF, SSMF lower concentrations of various immune indicators in the serum, although the differences were not marked. Significant differences were also observed in gut microbiota diversity and abundance between the two groups, particularly at the phylum and genus levels. Differential metabolic pathways were identified through KEGG pathway enrichment analysis, with ‘Metabolic pathways’ being the most significantly enriched. There were significant correlations between changes in certain gut microbiota and serum immune indicators, and most differential metabolites showed significant correlations with the gut microbiota. Research in this field will contribute to a deeper understanding of the mechanisms through which different rearing methods impact the health and metabolism of animals. In response to identified differences, we can delve into the biological mechanisms underlying them, subsequently formulating more rational livestock management strategies. This will provide essential scientific foundations for optimizing animal rearing practices, improving product quality, and promoting the sustainable development of the livestock industry.

## Author contribution

LY and AA were responsible for writing and revising the first draft and were involved in sample collection and processing. CP and HS were responsible for Conceptualization and design of experiments. SBM were responsible for project data processing and manuscript revision. YC, ZZ, JL, JZ and WZ are responsible for project management and fund preparation. All authors have read and agreed to the final manuscript.

## Ethics approval

Animal care and experiments were carried out in accordance with the criteria set by the Chinese Guidelines for the Care and Use of Laboratory Animals, and all methodologies were supervised and authorized by the Southwest University of Science and Technology’s Institutional Review Board.

